# An adaptive numerical method for multi–cellular simulations of tissue development

**DOI:** 10.1101/2024.01.06.574290

**Authors:** James M. Osborne

**Affiliations:** School of Mathematics and Statistics, University of Melbourne, Melbourne, 3010, Victoria, Australia

**Keywords:** Multi–Cellular Modelling, Cell–Based, Numerical Approximation, Adaptivity

## Abstract

In recent years, multi–cellular models, where cells are represented as individual interacting entities, are becoming ever popular. This has led to a proliferation of novel methods and simulation tools. The first aim of this paper is to review the numerical methods utilised by multi–cellular modelling tools and to demonstrate which numerical methods are appropriate for simulations of tissue and organ development and disease. The second aim is to introduce an adaptive time–stepping algorithm and to demonstrate it’s efficiency and accuracy. We focus on off–lattice, mechanics based, models where cell movement is defined by a series of first order ordinary differential equations, derived by assuming over–damped motion and balancing forces. We see that many numerical methods have been used, ranging from simple Forward Euler approaches through to higher order single–step methods like Runge–Kutta 4 and multi–step methods like Adams–Bashforth 2. Through a series of exemplar multi–cellular simulations, we see that if: care is taken to have events (births deaths and re–meshing/re–arrangements) occur on common time–steps; and boundaries are imposed on all sub–steps of numerical methods or implemented using forces, then all numerical methods can converge with the correct order. We introduce an adaptive time–stepping method and demonstrate that the best compromise between *L*_∞_ error and run–time is to use Runge–Kutta 4 with an increased time–step and moderate adaptivity. We see that a judicious choice of numerical method can speed the simulation up by a factor of 10–60 from the Forward Euler methods seen in Osborne *et. al*. [2017, https://doi.org/10.1371/journal.pcbi.1005387] and a further speed up by a factor of 4 can be achieved by using an adaptive time–step.

## 1 Introduction

Multi–cellular modelling, where tissues are modelled as a set of discrete interacting cells, is increasingly being used to study the development and malfunction of organs [19]. Multi– cellular modelling has been used to study: Liver development and disease [15]; Kidney development [9]; Skin development and wound healing [32, 33, 58]; Colorectal Crypt development and disease [16, 17, 54]; C. elegans Germline development [1]; Biofilm growth [47]; and viral immune system interaction within an infected tissue [45, 26], among others. These studies can be broken into two categories: studies following the growth and development of the tissue [1, 9, 45, 47]; or investigations into how perturbations to subcellular and cellular properties influence systems in dynamic equilibrium (where the properties of the system, i.e., numbers of cells, or average cellular velocity are approximately constant) [15, 16, 17, 32, 33, 54, 58].

Multi–cellular models can be broken up into two categories based on how cells are represented. The first of these is where cells are represented by objects that are restricted to a lattice, known as on–lattice models. The simplest example of on–lattice models are Cellular Automota (CA) models, where cells are represented as individual lattice sites (or collections of cells are an individual lattice site) [39]. Another example is the Cellular Potts Model (CPM) where cells are represented by collections of lattice sites [27]. Usually on–lattice models are simulated by defining a potential, *U*, of the system and using Monte Carlo simulation to evolve the system to minimise *U*. Stochasticity, intrinsic in the method, avoids the system being trapped at a local minimum. While being simple to implement and fast to run the constraint of being fixed to a lattice can introduce anisotropic effects in the simulations [39].

If the restraint of cells lying on a fixed lattice is removed then we get the second family of models known as off–lattice models. The simplest examples of off–lattice models are cell– center models like the Overlapping Spheres (OS) model (also known as a point based model) and Voronoi Tessellation (VT) model. In the former cells are represented as deformable spheres or as points with a decaying interaction force, and cells interact if they are within a specified interaction radius. In the latter cells are connected by a Delaunay Triangulation, with linear forces between cells, and cells are represented by the Voronoi Tessellation (of the cell centres) leading to polygonal cells [39]. Equations of motion are derived by balancing forces on cell centres. Additional off–lattice models can be created by having cells be represented by multiple components. The simplest of these is the Vertex Dynamics (VD) model where cells are represented as connected polygons whose vertices are free to move. Equations of motion are derived by considering a potential *U* and minimising with respect to moving vertices [21]. Each of these off–lattice models can be further specialised by how it defines free and interfacial boundaries and recent work has looked at the effect of this on simulation outcomes [23]. More detailed off–lattice models consider: arbitrary subcellular elements as point forces, the Subcellular Element Method (SEM) [38, 44]; arbitrary subcellular elements as finite elements, the Finite Element Model (FEM) [4]; or non connected cell boundaries, the Dynamic Cell Model (DCM) [8, 55].

In off–lattice models the motion of nodes (cell centres or components of cells) is represented by considering forces acting on the node, {**x**_*i*_}. These forces can be converted into cell motion by using Newtons second law which results in a second order equation of motion for each cell. Alternatively, these applied forces can be balanced by viscous drag forces, resisting motion of the nodes, by assuming that the motion is over–damped [39]. The assumption of over–damped motion leads to a system of *N* first order equations of motion, for the *N* nodes, of the form:

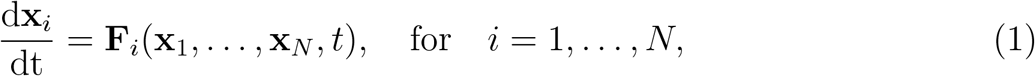

where **F**_*i*_ is the force exerted on node *i* (here the drag coefficient has been absorbed into the force for notational simplicity). The simplest approach for the solution of this system of equations, and therefore the simulation of off–lattice multi–cellular models, is to use a Forward Euler (FE) approximation to update the cell locations over time, [39]:

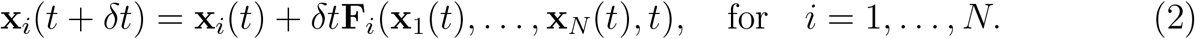

where *δt* is the chosen time–step. Usually a default time–step of around *δt* = 1*/*120≈ 2^−7^ or *δt* = 1*/*200 ≈ 2^−8^ is chosen, [39]. *δt* is chosen so that reducing it further doesn’t change the output of the simulation, to avoid oscillating numerical solutions i.e., to ensure that simulations are numerically stable.

Higher order numerical methods have also been used, for example higher order implicit schemes like Runge–Kutta methods [13] or multi–step schemes like Adams Bashforth 2 (AB2)[24]. However, there has been no detailed study into how these methods compare. An alternative simulation method which is used by some multi–cellular simulation tools is to use the energy formulation of the equations of motion and to use stochastic simulation, for example Metropolis sampling, to evolve the system [25, 30]. This has the advantage of speed and simplicity of implementation and can account for biological noise, however simulations are not 100% reproducible and it’s sometimes not clear if features of the simulation output are due to the model or due to the nature of the stochastic simulation process.

Over the years many multi–cellular computational tools have been developed. In Table 1 we present a list of popular off–lattice simulation tools, see Fletcher et. al. [19], or Vetter et. al. [55] for more complete lists. We choose not to include the popular off–lattice modelling tools EpiSim [50], Biocellion [28], and HAL [3], in our table, as they are Agent Based Modelling (ABM) frameworks which represent cells as off–lattice objects, however rather than being based on the equations of motion (or their potential form) movement is based on rules and stochastic simulation. We also don’t include popular on–lattice modelling frameworks like CC3D [49] and Morpheus [48] (CPM frameworks), or general ABM frameworks like Repast [7] and NetLogo [53].

**Table 1:**
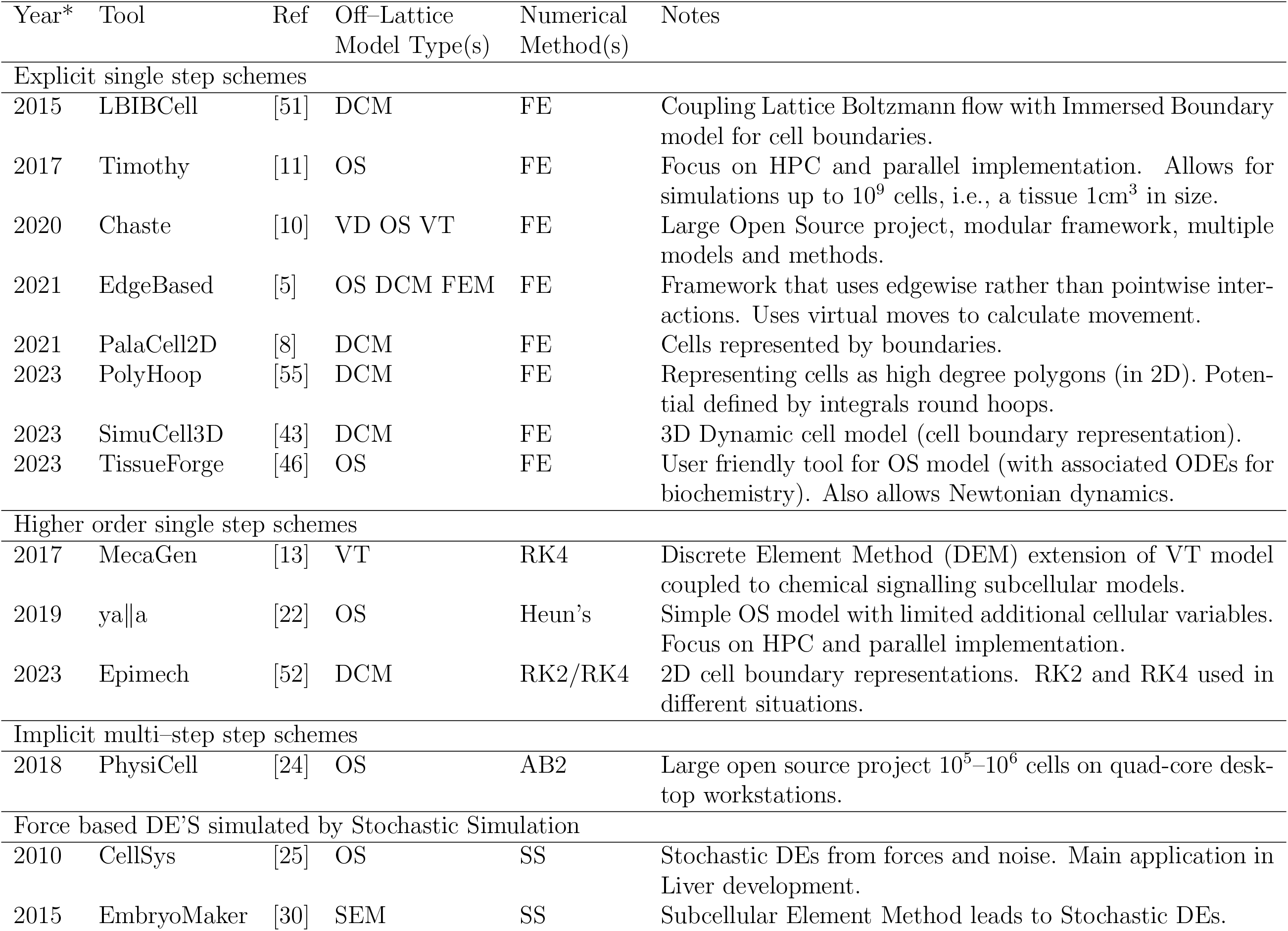
Off–lattice multi–cellular modelling tools. * denotes the year of the recent cited paper not the initial release. Methods legend: DCM–Dynamic Cell model; OS–Overlapping Speheres or Point based Force model; VD– Vertex Dynamics model; VT–Voronoi Tessellation; FEM–Finite Element Model; SEM– Subcellular Element Method; and ABM–Agent Based Model. Numerical methods legend: FE–Forward Euler (1st order Runge–Kutta) scheme; RK2–2nd order Runge–Kutta scheme; RK4–4th order Runge–Kutta scheme; Heun’s–Heun’s method (an alternative 2nd order Runge–Kutta scheme); AB2–2nd order Adams–Bashforth scheme; and SS–Stochastic Simulation.

Table 1 demonstrates the variety of multi–cellular models and simulation approaches in the literature. The most common numerical approach is FE as it is simple and well suited to the varying system sizes seen with multi–cellular models. Some frameworks use higher order methods (Heun’s method Runge–Kutta 4 (RK4) and AB2) where mechanical interactions are simple and speed is crucial [13, 22, 24]. Interestingly some tools have started to use different numerical methods for different conditions. For example EpiMech [52] uses a Runge–Kutta 2 (RK2) method for simulations without a substrate and a RK4 method for simulations where cells are attached to a substrate.

In other areas of mathematics, adaptivity, where time–step or spatial resolution is varied to ensure a chosen accuracy, have been used. These adaptive schemes can be based on analytic expressions or approximations for the error (for example in fluid flow [29]) or using a heuristic to inform the choice of numerical method or resolution (in cardiac electrophysiology [56, 57]). In either case a judicious choice of numerical method and the use of adaptivity can increase the accuracy of a solution method (for a given computational cost). It can reduce the computational cost by orders of magnitude while maintaining the same accuracy (ensuring stability).

In this paper we present a numerical framework for the simulation of off–lattice multi– cellular models which can be incorporated within existing simulation frameworks which use the over–damped assumption and a Forward Euler approximation [39]. The framework allows the use of single step methods such as Runge–Kutta (RK) schemes, and implicit schemes like Backward Euler (BE). This allows an increased accuracy for the same time– step. Therefore, allowing for larger time–steps (while maintaining the same accuracy) decreasing simulation runtime. In addition the numerical framework includes time–step adaptivity which allows the maintenance of accuracy while decreasing runtimes further.

The remainder of this paper is structured as follows. Firstly, in Section 2, we present the multi–cellular models and numerical methods used in this study, along with a set of exemplar multi–cellular simulation problems. Secondly, in Section 3, we present simulations of the exemplar problems using the numerical methods presented and demonstrate the convergence properties for each exemplar, we also demonstrate the power of using an adaptive time–step. Finally, in Section 4, we discuss balancing accuracy and runtime (including with adaptivity) along with the main convergence results and present some avenues for future work.

## 2 Methods

In this section we present: the off–lattice multi–cellular models (Section 2.1); the numerical methods (Section 2.2); and the exemplar multi–cellular problems (Section 2.3), which are used in this study.

### 2.1 Off–lattice multi–cellular models

As discussed in the introduction off–lattice models consider the evolution of cells within a tissue by tracking the motion of “nodes”, **x**_1_, …, **x**_*N*_ ∈ℛ^*n*^ (*n* = 1, 2, 3), which represent cell centres, subcellular components, or components of the cell boundary. Here we consider the sub–class of off–lattice models where motion is assumed to be over–damped. So the equations of motion for each node, **x**_*i*_, reduce to a force balance [39]

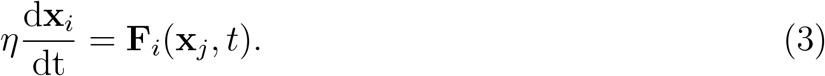

where **F**_*i*_ is the force applied to the “node” which is balanced by the viscous drag (drag coefficient, *η*) experienced. We describe three off–lattice models: overlapping spheres, Voronoi tessellation; and vertex dynamics. Here, we restrict ourselves to the model components that are relevant for our study, and include mainly a high level description. Note that for simplicity all cells in this study have identical properties and we also exclude the growth phase in each model (i.e., all cells are mature). All results presented here also hold for heterogeneous tissues and for growing cells. More detailed model descriptions, including extensions to have cell growth and homogeneous cells can be found in Osborne et. al. [39] or Germano et. al. [23]. All model parameters used in this study are given in Table 2.

**Table 2:** Mechanical parameter values used in all exemplar simulations. Parameter values sourced from [23, 39]. Note, CD=Cell Diameters CM=Cell Mass.

| Parameter | Description | Model | Value | Units |
| --- | --- | --- | --- | --- |
| $\eta$ | Drag coefficient | All | 1.0 | CM hours<sup>-1</sup> |
| $\mu$ | Elastic interaction constant | OS | 50.0 | CM hours<sup>-2</sup> |
| $\rho$ | Decay in attractive force | OS | 5.0 | - |
| $r_{\max}$ | Force cut-off length | OS | 1.5 | CD |
| $s_{ij}$ | Mature cell separations | OS & VT | 1.0 | CD |
| $\kappa$ | Spring constant | VT | 50.0 | CM hours<sup>-2</sup> |
| $\alpha$ | Area deformation coefficient | VD | 50.0 | CM hours<sup>-2</sup> CD<sup>-2</sup> |
| $\beta$ | Membrane tension coefficient | VD | 1.0 | CM hours<sup>-2</sup> |
| $A^T$ | Cell target area | VD | 1.0 | CD<sup>2</sup> |
| $\gamma_{\text{cell,cell}}$ | Cell-cell adhesion coefficient | VD | 1.0 | CM CD hours<sup>-2</sup> |
| $\gamma_{\text{cell,boundary}}$ | Cell-boundary adhesion coefficient | VD | 2.0 | CM CD hours<sup>-2</sup> |
| $d_r$ | Cell rearrangement threshold | VD | 0.01 | CD |

### Overlapping Spheres (OS) model

Sometimes referred to as point force model. Cells are represented by their centres, **x**_*i*_ ∈ ℛ^*n*^ (*n* = 1, 2, 3), which are points free to move in space. These models were first used by [14], who defined the framework but used energy minimisation to simulate cell movement. later works [40] extended the model and moved to used the simulation methods illustrated here. 2D and 3D examples can be seen in Figure 1 (b) top and bottom row. Cell connectivity is defined by finding all cells within a specific interaction radius, 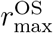. Cell connectivity changes as cells move in or out of this interaction radius. The equation of motion for cell centre *i*, **x**_*i*_, is given by, [39],

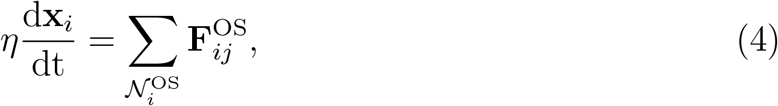

where *η* is the drag coefficient (which could vary for each cell but here is constant). 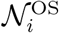 is the set of all neighbours of cell *i*, defined as those closer than 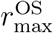, and 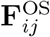 is the force exerted on cell *i* by cell *j* which is given by

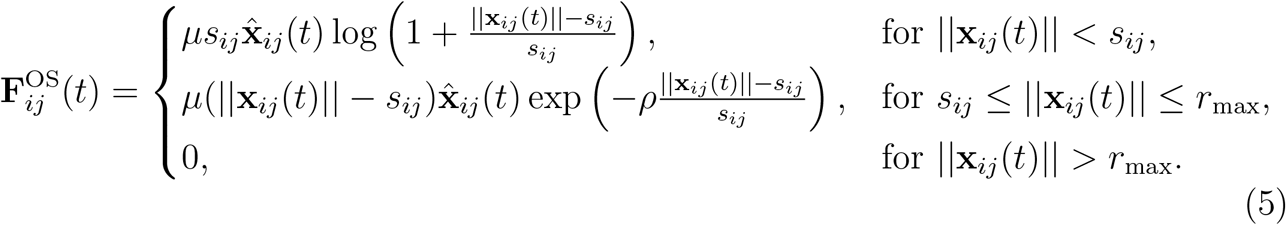

**Figure 1.**
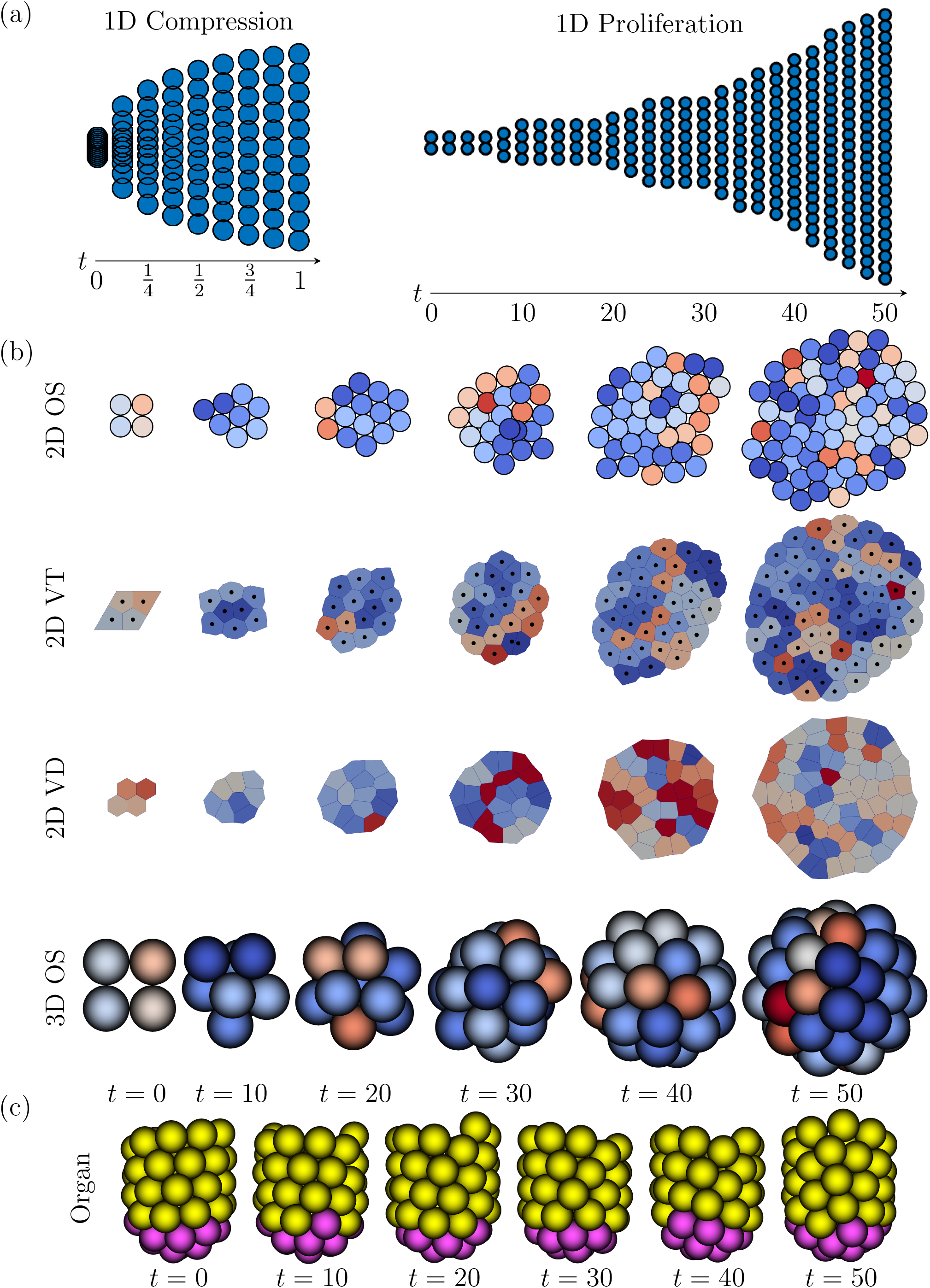
Exemplar problems. Time series snapshots from the chosen exemplar problems. (a) 1D Compression and 1D Proliferation exemplars. (b) 2D Monolayer and 3D Spheroid exemplars. (c) 3D Organ exemplar.

Here *μ* and *ρ* are parameters that control the magnitude of the force and the decay of the attractive force respectively, which could depend on cell properties but here are constant. **x**_*ij*_ (*t*) = **x**_*i*_(*t*) − **x**_*j*_ (*t*), and 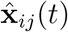 is the corresponding unit vector. *s*_*ij*_ is the natural separation between cells *i* and *j* which is the sum of the radii, and here all cells have a radius of 0.5CD (Cell Diameters) so *s*_*ij*_ = 1CD. When a cell divides a random direction is chosen and 2 child cells are placed apart by 1CD. More details and a full model description/derivation can be found in [1, 41].

#### Voronoi Tessellation (VT) model

The Voronoi Tessellation model was originally proposed in [31] for studying the colonic crypt but has been extended for use in various other epithelial tissues [39]. Again cells are represented by their centres, **x**_*i*_ ∈ *ℜ*^*n*^ (*n* = 1, 2, 3), which are free to move, but here cell connectivity is defined by the Delaunay Triangulation and cell shape is defined by the Voronoi Tessellation [39]. This connectivity needs to be re–calculated regularly as cells move and cell connections change. 1D and 2D example simulations can be seen in Figures 1 (a) and (b) second row. Cell motion is again calculated by balancing the forces on the cell centres but here there is a linear spring between connected cell centres, [39]

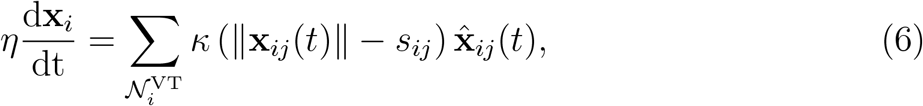

Where *η* is again the drag coefficient (which could vary for each cell but here is constant). 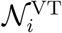 is the set of all neighbours of cell *i* which here is calculated from the Delaunay Triangulation. *κ* is the spring constant and controls the magnitude of the force, which could depend on cell properties but here is constant. **x**_*ij*_(*t*) and 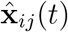 are as for the OS model. *s*_*ij*_ is the target length of the edge between cells *i* and *j* which here is 1CD for all cells. When a cell divides a random direction is chosen and 2 child cells are placed apart by 0.1CD and connectivity is recalculated. In order to visualise we use the bounded VT model from [23].

#### Vertex Dynamics (VD) model

Here we restrict ourselves to the 2 dimensional version of the model, as presented in [20, 21], originally developed in [35, 36, 37]. Here cells are represented by polygons whose vertices, **x**_*i*_ ∈ *ℜ*^2^, are free to move in space. Cells are connected if they share a vertex. Cell connectivity is updated by a process known as T1 swaps which is a local re–meshing operation which allows cells to move past each other and occurs when edges are less than *d*_*r*_ in length, see [20] for details. A similar class of VD models, using the same structure but with different forces, are defined in [18], and the convergence results presented here still hold. An example simulation can be seen in Figure 1 (b) third row. Cell motion is calculated by balancing forces on the cell vertices by minimising an energy functional, containing terms to keep cells approximately circular with a given target area.

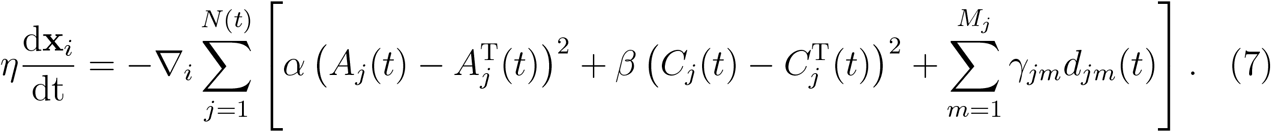

Where *η* is again the drag coefficient (which could vary for each vertex but here is constant). 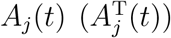 and 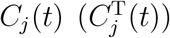 are the area (target) and perimeter (target) of cell *j* at time *t*, respectively. Here we assume that every cell has the same constant target area, 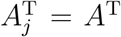, and perimeter, 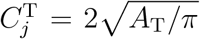. Note, usually the VD model has cell growth over G1 phase of the cell cycle to reach the mature target area but we used a fixed size, of *A*_T_ = 1, for simplicity, the results presented here still hold for other target areas or growing cells. *α* and *β* describe cell compressibility and cytoskeletal tension, respectively, *M*_*j*_ denotes the number of vertices contained in cell *j, d*_*jm*_(*t*) is the length of edge *m* in cell *j* and *γ*_*jm*_ describes cell adhesion in edge *m*, which depends on whether the edge is on the boundary, or is internal to the tissue,

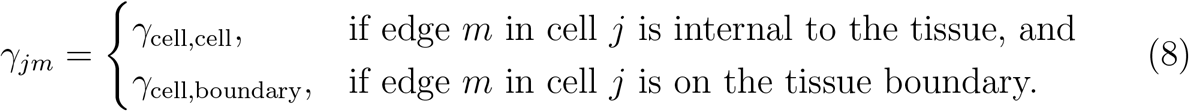

Force originally from [35]. When a cell divides it does so by calculating the long axis and adding new vertices to make two child cells [20].

### 2.2 Numerical Methods

For each model we are solving Equation (4), (6), or (7) which can all be written as the system

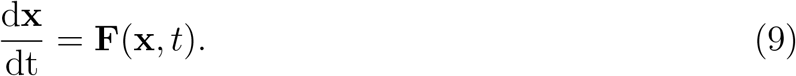

where **x** = (**x**_1_, … **x**_*N*_) are the locations of cell centres (OS and VT) or cell vertices (VD), and **F**(**x**, *t*) represents the force applied (and includes the drag coefficient, *η*), which is model dependent. In order to solve this we generate a numerical approximation to this system by introducing a series of *N*_S_ time–steps

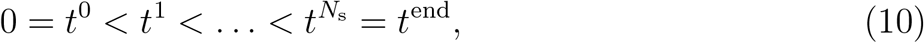

where *t*^*i*^ = *i* × *δt* and *δt* = *t*^end^*/N*_S_ is the time–step. We then use a numerical method to calculate the sequence

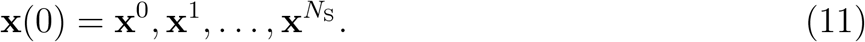

which gives us an approximation to the solution to the problem at the chosen time–steps.

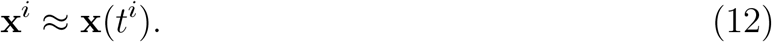

This sequence can be calculated by many different methods. The most popular is the Forward Euler (FE) method which calculates this sequence using.

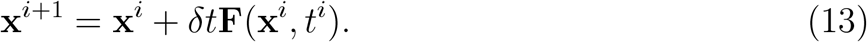

In this study we focus on single step methods using the setup above. We use both explicit, where we end up with an explicit formula for the next term of the sequence (as for FE), and implicit methods, where we end up with an implicit formula for the next term in the sequence.

#### Explicit Methods

The most common collection of explicit single step numerical methods are known as Runge– Kutta methods [6]. In these methods we discretise the derivative as for Forward Euler, however, we also evaluate **F**(**x**, *t*) at a collection of intermediary points instead of just at **x**^*i*^ and *t*^*i*^. This gives rise to a set of updates.

#### Forward Euler (FE) method

The update for FE is given by Equation (13). This method has a theoretical order of convergence of 1 so as you halve *δt* you should halve the global truncation error (the error of the numerical approximation to the displacements compared to the exact solution).

##### MidPoint (MP) method

The update for the MP method (also known as Runge–Kutta 2) is, [6],

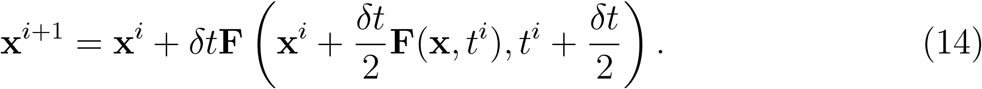

This method has a theoretical order of convergence of 2 so as you halve *δt* you should reduce the global truncation error by a factor of 4.

##### Runge–Kutta 3 (RK3) method

The update for the RK3 method is, [6],

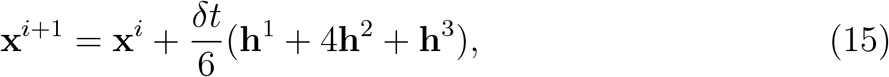

where

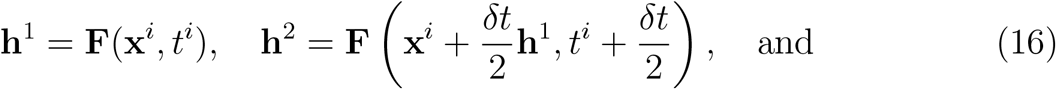

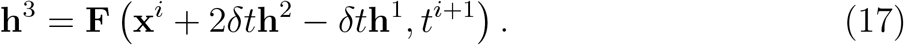

This method has a theoretical order of convergence of 3 so as you halve *δt* you should reduce the global truncation error by a factor of 8.

##### Runge–Kutta 4 (RK4) method

The update for the RK4 method is, [6],

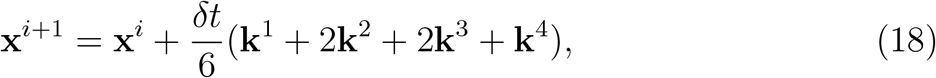

where

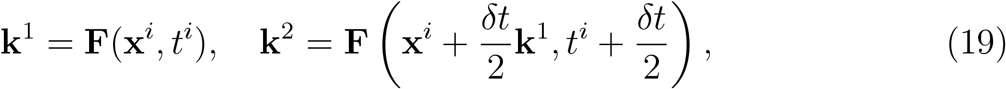

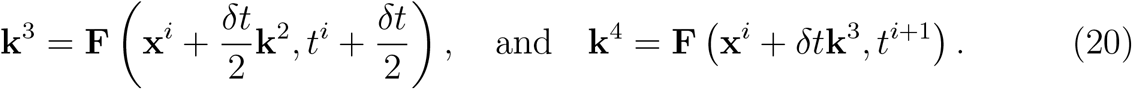

This method has a theoretical order of convergence of 4 so as you halve *δt* you should reduce the global truncation error by a factor of 16.

#### Implicit Methods

An alternative to these explicit Runge–Kutta schemes is to generate an implicit equation for **x**^*i*+1^. This can lead to increased stability. i.e., the solution may not diverge for larger *δt* as can be the case for explicit methods [6]. However, rather than writing the update **x**^*i*+1^ explicitly we have an implicit equation to solve for **x**^*i*+1^, which can be costly.

##### Backward Euler (BE) method

The update for the BE method is [6]

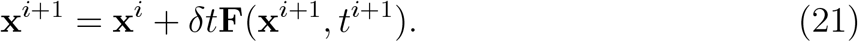

This method has a theoretical order of convergence of 1 so as you halve *δt* you should halve the global truncation error.

##### Adams Moulton (AM) method

The update for the second order AM method (also known as the Trapezium Rule) is [6]

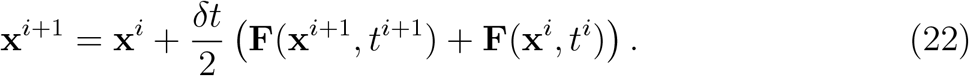

This method has a theoretical order of convergence of 2 so as you halve *δt* you should reduce the global truncation error by a factor of 4.

In order to solve Equations (21) and (22) for **x**^*i*+1^ we need to resort to numerical root finding methods. Here we use the Newton Method based non–linear solvers from PETSc [2] with a relative and absolute tolerance of *δ*_TOL_. Note that higher order AM methods don’t fit within the framework presented in this paper as they are multi–step methods i.e., they use solutions from the previous 2 (or more) time–steps to calculate **x**^*i*+1^.

#### Adaptivity

An adaptive time–step is a way to use a smaller time–step when the solution is varying rapidly and a larger one when the solution is varying more slowly. For some systems an approximation to the global truncation error can be calculated from the problem (*a priori* error analysis) or the problem and solution (*a posteriori* error analysis) to allow an appropriate time–step to be calculated [6]. However, due to the changing number and form of the system of equations which comprise, multi–cellular models such analysis is not feasible so instead we use a heuristic error analysis by introducing an Absolute Movement Threshold, *x*_AMT_, which defines a maximum distance a cell can move in one time–step, and use this to inform an adaptive time–step 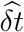.

The way we implement adaptivity is that if a proposed update, **x**^*i*+1^ causes nodes to move more than the threshold, specifically max {Δ**x**^*i*^} = max {|**x**^*i*+1^ − **x**^*i*^|} *> x*_AMT_, (where the maximum is taken over the set of all nodes). then the time–step is reduced by a power of 2 (i.e., successively halved) in order to ensure max {Δ**x**^*i*^} ≤ *x*_AMT_. The adaptive time–step 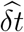 is kept at this level until we reach the end of the global time–step, *δt* where the time–step is reset to *δt*. Full details of the adaptive time–step algorithm are given in Algorithm 1.

The numerical parameter values used in this study are given in Table 3. Note that we use a range *δt* = [2^−16^, 2^−6^] ≈ [1.5 × 10^−5^, 1.6 × 10^−2^] for all exemplars presented in this study. This allows us to capture the convergence, or not, of all methods without using values of *δt* with infeasibly long run times (greater than 10 hours).

**Table 3:** Numerical parameter values used in all exemplar simulations. Note, CD=Cell Diameters.

| Parameter | Description | Value | Units |
| --- | --- | --- | --- |
| $\delta t$ | Time-step | $2^{-6}, 2^{-8}, \dots, 2^{-18}$ | hours |
| $x_{\text{AMT}}$ | Movement threshold | $1, 0.1, \dots, 10^{-5}$ | CD |
| $\delta_{\text{TOL}}$ | Non-linear solver tolerance | $10^{-5}, 10^{-10}$ | CD |

##### Algorithm 1

**Simple Adaptivity Algorithm**. Time–step is reduced to prevent movements of nodes larger than the threshold *x*_AMT_. The time–step, 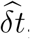, is reset to the base value, *δt* every global time–step.

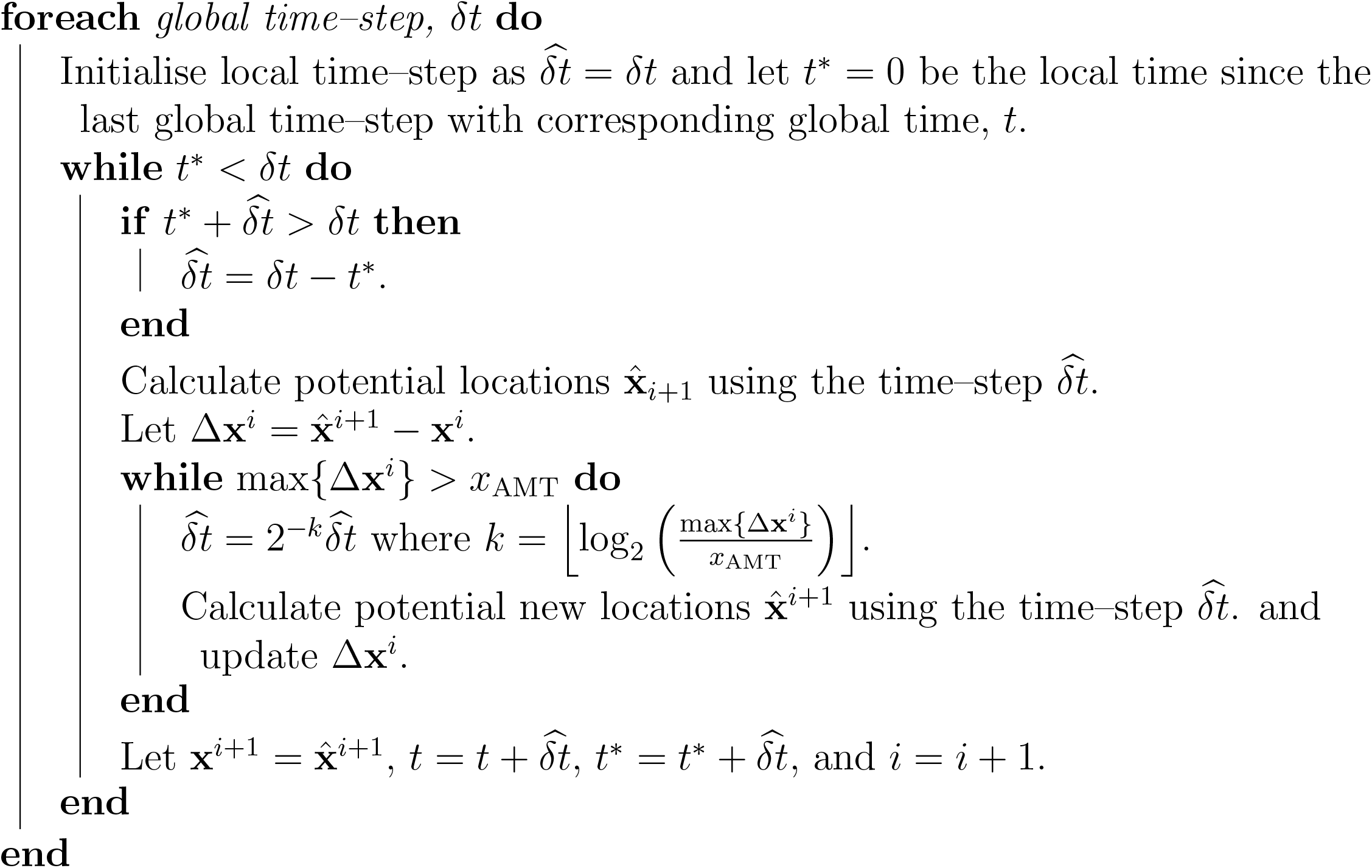

### 2.3 Exemplar problems

In order to demonstrate the usage, and compare the efficiency, of the numerical methods presented above, when applied to multi–cellular simulations, we use a set of exemplar problems that increase in complexity. The exemplar problems provide a surrogate for multi–cellular simulations used in the literature (both growing domains and systems in homeostasis) are presented in Figure 1 and detailed below.

#### 1D Compression

The simplest multi–cellular simulation is the relaxation of a chain of cells under compression. Motivated by [12, 41], we track the position of 10 cell centers connected by springs (VT model) initially compressed to 10% of their size and allow the system to relax for 1 hour. A simulation of this exemplar can be seen in Figure 1 (a). We can compare the trajectories calculated from numerical approximation against the analytic solution **x**^Exact^(*t*) (see Supplementary Section Appendix B for details of how to calculate the analytic solution) by calculating the *L*_∞_ error over time, *L*_∞_(*t*^*i*^) = max {|**x**^*i*^− **x**^Exact^(*t*^*i*^) |}. This exemplar allows us to demonstrate the theoretical orders of convergence of the chosen numerical methods in a system with a fixed number of equations (cells).

#### 1D Proliferation

The second exemplar we consider is the deformation of a chain of proliferating cells. Motivated by [34], we consider a proliferating 1D chain of cells (VT model). We start with two cells (placed 1CD apart) which divide once they reach a given age (drawn on cell birth from a *U* (7, 17) distribution (i.e., mean cell cycle duration is 12 hours. We run the simulation for 50 hours. A simulation of this exemplar can be seen in Figure 1 (a). We use the same 1D linear spring model as in the compression example and compare simulations against exact solution using extracted division events (details in Supplementary Section Appendix B) and calculate the *L*_∞_ error over time. This exemplar allows us to demonstrate how the numerical numerical methods perform when the number of equations increases over time due to changing cell number.

#### 2D Monolayer

The next exemplar is following the movement of cells within a monolayer of proliferating cells. Motivated by [23, 39], we look at a growing monolayer of cells in 2D using the OS, VT, and VD models. We start with 4 cells placed in equilibrium as shown in Figure 1 (b) and allow cells to proliferate as in the 1D Proliferation exemplar. The simulation is then run for 50 hours and the trajectories tracked, see Figure 1 (b). We compare simulations to the run using RK4 with *δt* = 2^−18^ and calculate the *L*_∞_ error over time, 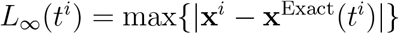. Note, this value of *δt* is sufficiently small so that using a smaller value doesn’t change any of the convergence plots presented. This exemplar allows us to demonstrate the how the numerical numerical methods are affected by cell re–arrangements as the tissue grows.

#### 3D Spheroid

We extend the 2D monolayer exemplar to 3D by looking at the movement of cells within a spheroid of proliferating cells using an OS model. We again start with 4 cells placed in equilibrium as shown in Figure 1 (b), bottom row, and allow cells to proliferate as before. The simulation is then run for 50 hours and the trajectories tracked, allowing the *L*_∞_ error over time to be calculated in the same way as the 2D monolayer exemplar. This exemplar allows us to demonstrate the how the numerical methods behave on 3D tissues.

#### 3D Organ

Our final exemplar looks at how cell turnover drives cell movement on a fixed domain. Motivated by simulations of colorectal crypts from [17] we use a 3D OS model and constrain cells to lie on a test tube shaped geometry (base radius 1.5CD and height 4CD) using a restraining force (which leads to cells lying on a fixed surface), see Figure 1 (c). We divide the domain into a proliferative region for *z* ≤ 2CD (pink cells) where cell division occurs as in the 3D Spheroid exemplar, and a differentiated region *z >* 2CD (yellow cells) where cells don’t proliferate (position dependent cell proliferation). Cells are removed when they reach the top of the domain, *z* = 4CD (position dependent cell removal). To initialise simulations we start with 4 cells at the base of crypt and run for 100 hours (with *δt* = 2^−6^ using RK4) to get to a developed crypt in dynamic equilibrium (see Figure 1 (c)). We then use this as the initial condition and simulate cell movement for a further 50 hours. We compare simulations to the run using RK4 with *δt* = 2^−18^ and calculating the *L*_∞_ error over time, 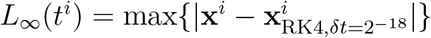. Note, this value of *δt* is sufficiently small so that using a smaller value doesn’t change any of the convergence plots presented. This exemplar allows us to demonstrate the how the numerical numerical methods behave when simulating a 3D homeostatic tissue.

#### 2.4 Implementation

All simulation were undertaken using the Chaste simulation framework. All code used is available, as a bolt on user project, under an Open Source licence, at https://github.com/jmosborne/MultiCellularNumericalMethods.

## 3 Results

To compare the performance of the chosen numerical schemes we simulate our exemplar systems for varying *δt* = 2^−*k*^ (for *k* = 6, 8, …, 16) and compare the results. In order to make numerical convergence possible we make the following assumptions with all simulations presented in this study:

1. all birth or death events occur only at the largest time–step, *δt* = 2^−6^;
2. all re–meshing events: recalculation of Voronoi Tessellation (VT); undertaking T1 swaps (VD); or recalculation of neighbours (OS), only occur at the largest time–step, *δt* = 2^−6^; and
3. all random events (birth times and division angles) are seeded so they are the same for each simulation run.

We now look at each exemplar problem in turn and discuss how the numerical methods presented perform.

### 3.1 Comparison with exact solution shows expected numerical convergence when events are synchronised

Figure 2 shows the *L*_∞_ error over time, the convergence of the *L*_∞_ error and the dependence of the *L*_∞_ error on computational runtime, *T*_R_, for the 1D Compression ((a) and (b)) and 1D Proliferation ((c) and (d)) exemplars. We see that the error is approximately constant over time for the 1D Compression exemplar, and constant between division events for the 1D Proliferation exemplar. In both cases the *L*_∞_ error converges as per the theoretical convergence rates (e.g. halving *δt* reduces *L*_∞_ error by a factor of 16, demonstrating order 4 convergence for RK4) see Figure 2. Note that in the 1D Proliferation exemplar if cells proliferate on non common time–steps then only first order convergence is witnessed in all methods (results not included for brevity).

**Figure 2.**
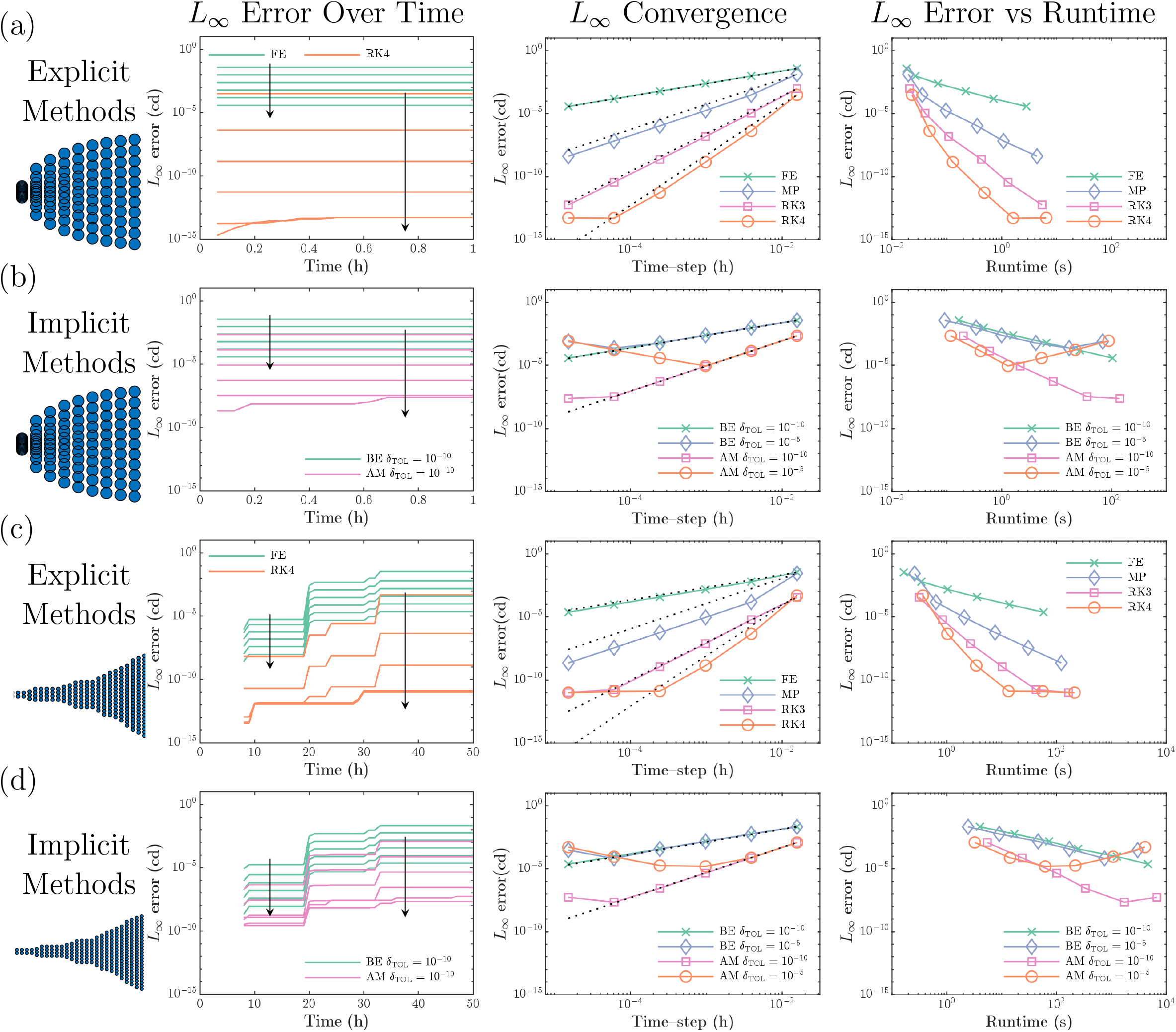
One dimensional chain of cells. (a) and (b) one dimensional chain of cells relaxing from compression over an hour (1D Compression exemplar), simulated using explicit (a) and implicit numerical methods (b). (c) and (d) one dimensional chain of cells with proliferation (mean cell cycle duration 12 hours) up to 50 hours (1D Proliferation exemplar), simulated using explicit (b) and implicit numerical methods (d). For explicit methods we present the L _∞_ error over time for both the FE method (green) and RK4 method (red). For implicit methods we present the L_∞_ error over time for both the BE method (green) and AM method (purple) (for low numerical tolerances *δ*_TOL_ = 10^−10^). We also show how the maximum of this L_∞_ error varies with time{step (*δt*) and runtime (T_R_) for all numerical schemes considered here. Arrows show the direction of decreasing *δt*. Dotted black lines in the L_∞_ convergence plots show the theoretical order of convergence for each numerical method. Note in all cases simulation runtimes scale linearly with *δt* see Supplementary Figure S1.

Figures 2 (b) and (d) show that the convergence for implicit methods is affected by the convergence of the root finding algorithms. As the *L*_∞_ error of the solution decreases the error of the root finding algorithm cancels out the increases in accuracy. This is more prevalent in the case of a higher solver tolerance (*δ*_TOL_ = 10^−5^, Figure 2 (b)). While the implicit methods converge as expected for low solver tolerance (*δ*_TOL_ = 10^−10^) they take much more computational time over the explicit methods (See Figure 2 right column and Supplementary Figure S1) therefore we choose to focus on the explicit methods moving forward.

We note that for both the 1D exemplars the solution using RK4 with *δt* ≤ 10^−16^ is equal to the exact solution to machine precision (taken to be *<* 10^−12^), Figures 2 (a) and (c). From now on we will use the RK4 approximation with 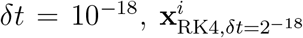, as the “exact solution”. We also note that the convergence behaviour of the explicit methods is illustrated well by first order convergence of the FE method and the fourth order convergence of the RK4 method, therefore we will focus on these 2 methods moving forward.

### 3.2 Dynamic tissue connectivity can affect convergence and accuracy in 2D and 3D

In order see how cell rearrangement influences the performance of the numerical methods we simulate the 2D Monolayer (for VT, VD, and OS models) and 3D Spheroid (OS model) exemplars for decreasing *δt* and compare the results to the “exact solution”^1^,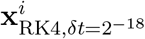.

Figure 3 (a)–(c) and (d) shows the *L*_∞_ error over time, the convergence of the *L*_∞_ error and the dependence of the *L*_∞_ error on computational runtime, for the 2D Monolayer and 3D Spheroid exemplars, respectively. We see that for the 2D Monolayer exemplar we maintain convergence rates seen in 1D, Figures 3 (a)–(c), this convergence is seen after a delay (to *δt* = 2^−8^) in the VD model. By looking at the *L*_∞_ error over time we see that the error increases due to the occurrence of an increased number of cells and division and “re–meshing” events. In reality, when using simulations like these are used, multiple runs are made (for different random seeds) and averages presented. Note that, if we allow re–meshing events (or divisions) to occur on any time–step we reduce down to first order convergence for all methods (results not shown for brevity).

**Figure 3.**
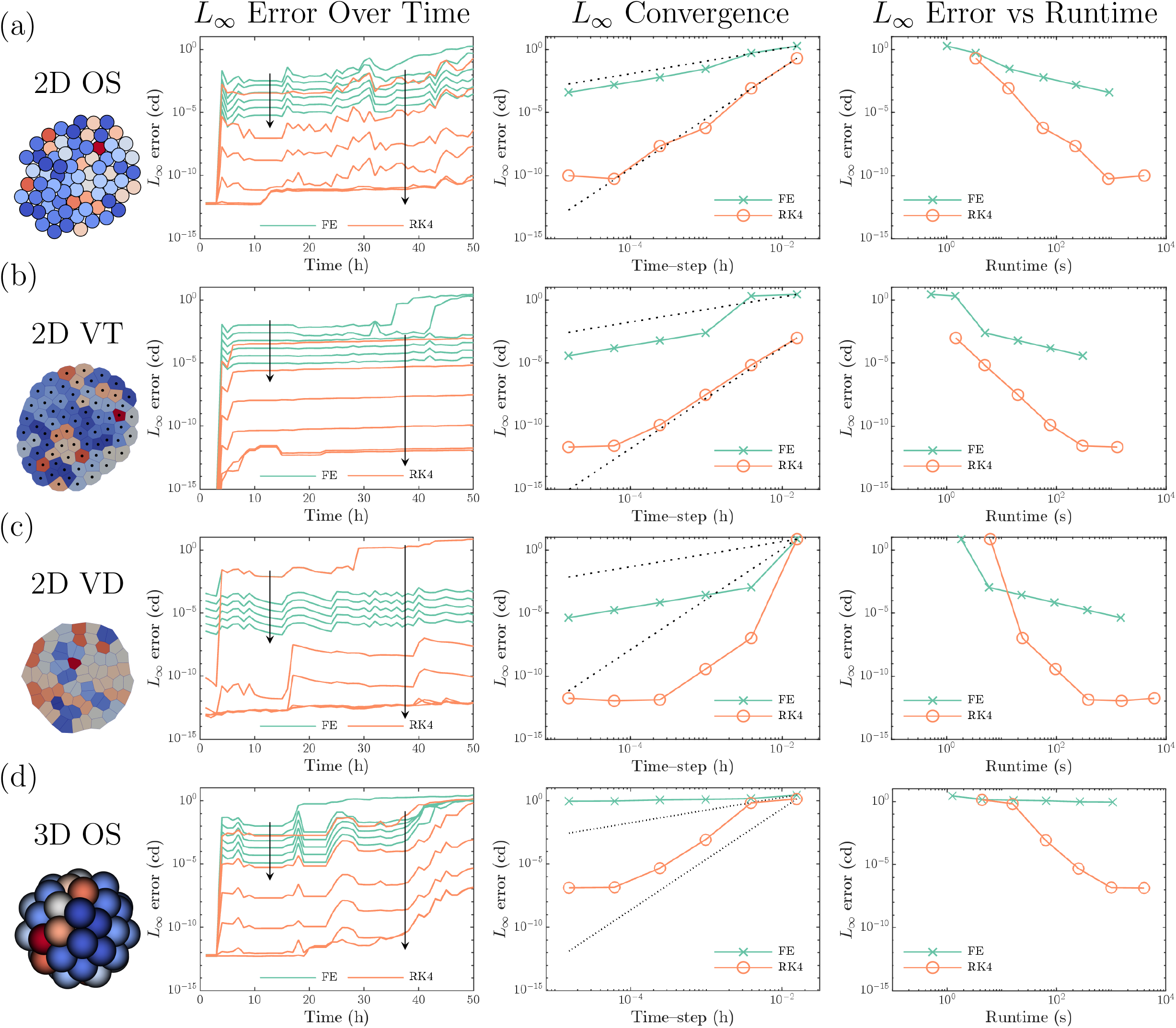
Growing monolayer and spheroid. Simulations of tissues with 4 initial cells and stochastic proliferation (mean cell cycle duration 12 hours) up to 50 hours (2D Monolayer and 3D Spheroid exemplars). For each simulation type, (a) OS model in 2D (b) VT model in 2D, (b) VD model in 2D, and (d) OS model in 3D. For each model we present the L_∞_ error over time for both the FE method (green) and RK4 method (red), we show how the maximum of this L_∞_ error varies with time-step *(δ*t) and runtime (T_R_). Arrows show the direction of decreasing *δt*. Dotted black lines in the L_∞_ convergence plots show the theoretical order of convergence for each numerical method. Note simulation runtimes scale linearly with *δt* see Supplementary Figure S2.

We see that the FE method is unable to converge for the 3D Spheroid exemplar, Figure 3 (d). This is because in this exemplar there is more movement and cell rearrangement than the 2D Monolayer exemplar, which increases the error (shown in Figure 3 (d), left). This increase also occurs in the RK4 simulations however, the RK4 method is still able to converge, giving order 4 convergence down to an *L*_∞_ error of 10^−8^.

All RK4 simulations (for *δt* ≤ 2^−8^) have a lower *L*_∞_ error than FE simulations of a similar runtime, Figure 3 right column. This shows that the same error can be achieved using RK4 with fewer time–steps and a quicker runtime than using FE. Looking at the the most accurate FE simulation (where FE is still performing well, i.e., in 2D simulations) we see that the RK4 simulation with the same *L*_∞_ error are all over an order of magnitude faster than the equivalent FE simulations, Figure 3 (a)–(c) right column. For 3D simulations all RK4 simulations (*δt <* 2^−8^) exhibit a lower *L*_∞_ error than is possible using the FE method for any *δt* used here, Figure 3 (d). Therefore we see that using an RK4 method with appropriately chosen *δt* can achieve the same *L*_∞_ error as a FE simulation over 10 times faster.

### 3.3 Convergence during dynamic equilibrium requires successively smaller time–steps

In Figure 4 we present simulations of our 3D Organ exemplar, in (b) we show the *L*_∞_ error over time (compared to the “exact solution”, 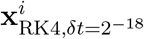) for the FE and RK4 methods, and in (c) we present the convergence of the *L*_∞_ error, as we reduce the time–step, after set simulation times (*t* = 10, 25, and 50 hours).

**Figure 4.**
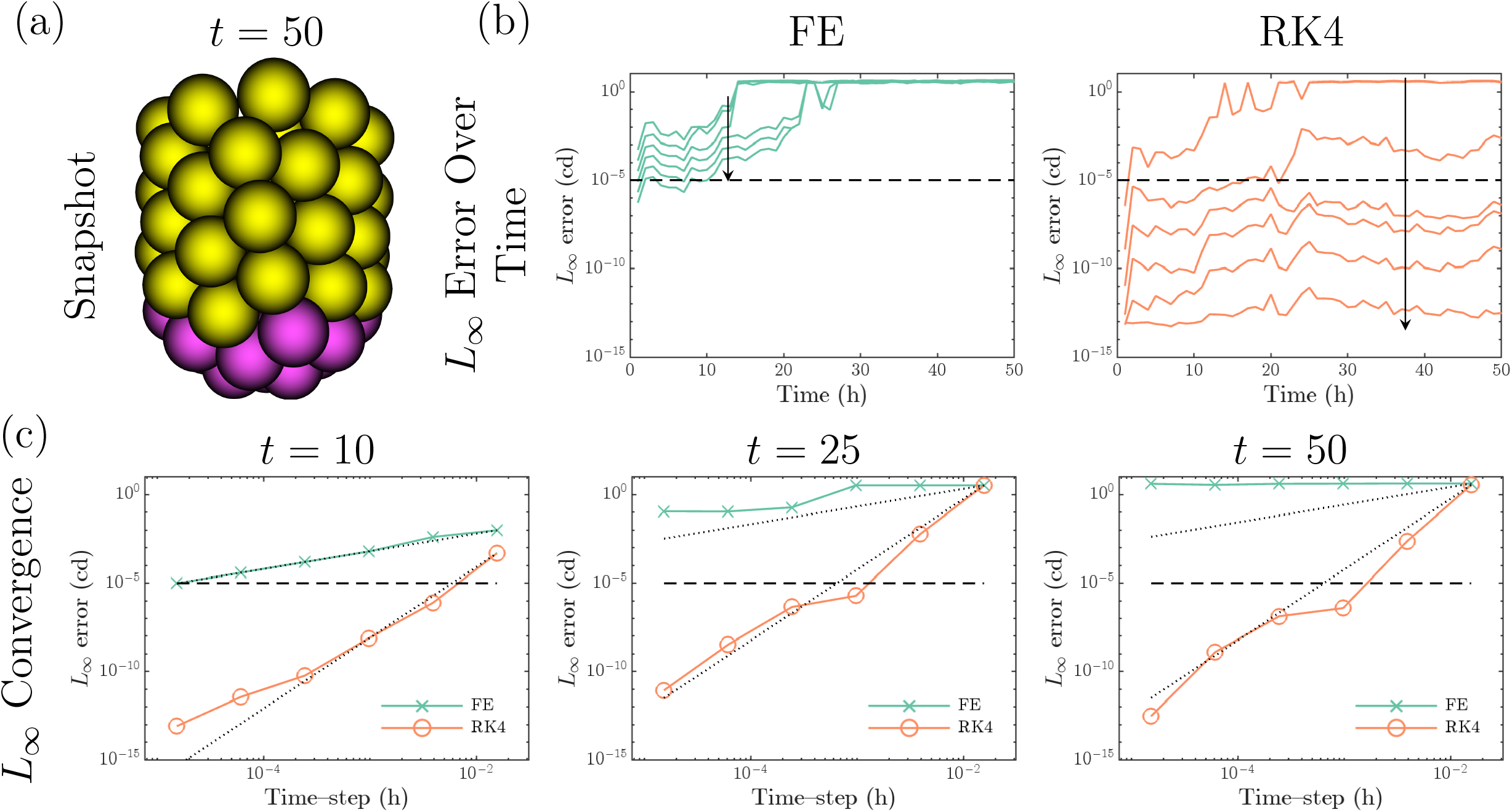
Convergence for system in dynamic equilibrium. Simulation of organ system with boundary conditions, and position dependent cell proliferation and removal. (a) example of simulation in dynamic equilibrium (3D Organ exemplar). (b) L_∞_ error over time for both the FE method (green) and RK4 method (red). Arrows show the direction of decreasing *δt*. (c) convergence of L_∞_ error at t = 10; 25; and 50 hours for both the FE method (green) and RK4 method (red). Dotted black lines show the theoretical order of convergence for each numerical method. In (b) and (c) the target L_∞_ error of L_∞_ = 10^−5^ is shown by the black dashed line.

In contrast to the 3D spheroid exemplar, we observe from Figure 4 (b) that if the *L*_∞_ error is below 10^−5^ at *t* = 25 then it stays below 10^−5^ at *t* = 50. Therefore, we consider a target *L*_∞_ error of 10^−5^ (to denote stability of numerical method) and this is shown by the black dashed line in Figures 4 (b) and (c). We also see that FE is not able to maintain a low *L*_∞_ error for any time–step used here (*δt* ≥2^−16^), Figure 4 (b). FE could be able to maintain a low error for some *δt* ≤ 2^−18^, however these simulations would take at least 2 ×10^4^ seconds ≈ 5.5 hours to run so aren’t feasible. Our choice of using forces to constrain cells to a fixed geometry gives more reliable convergence than simpler alternatives. Results with hard boundaries (as implemented in [17]) lose convergence faster than the force based results presented here (results not shown for brevity).

### 3.4 Adaptive calculation of time–step gives mechanism to balance speed and accuracy

From Figure 4 we see a time–step of *δt* = 2^−10^ (for RK4) is necessary to ensure that the *L*_∞_ error remains below the target of *L*_∞_ = 10^−5^ (and that it’s not possible for FE). In Figure 5 (a) we compare the convergence of the *L*_∞_ error as we include increased levels of adaptivity, *x*_AMT_ = 1, 0.1, 0.01, …, 10^−5^. In Figures 5 (b) and (c) we present the *L*_∞_ error and runtime for RK4 simulations (FE versions are in Supplementary Figure S3). We can see that decreasing *x*_AMT_ leads to a subsequent decrease in *L*_∞_ error and this is maintained over time for the RK4 method even for larger global time–steps, *δt*. For the FE method adaptivity improves the maintenance of convergence however the only case that maintains a low *L*_∞_ error (*L*_∞_ ≈ 10^−4^) for *t* = 50 hours, is for *δt* = 2^−16^ and *x*_AMT_ = 10^−5^ which is computationally prohibitive (runtime is over 10^4^ seconds, shown in Supplementary Figure S3 (c)).

**Figure 5.**
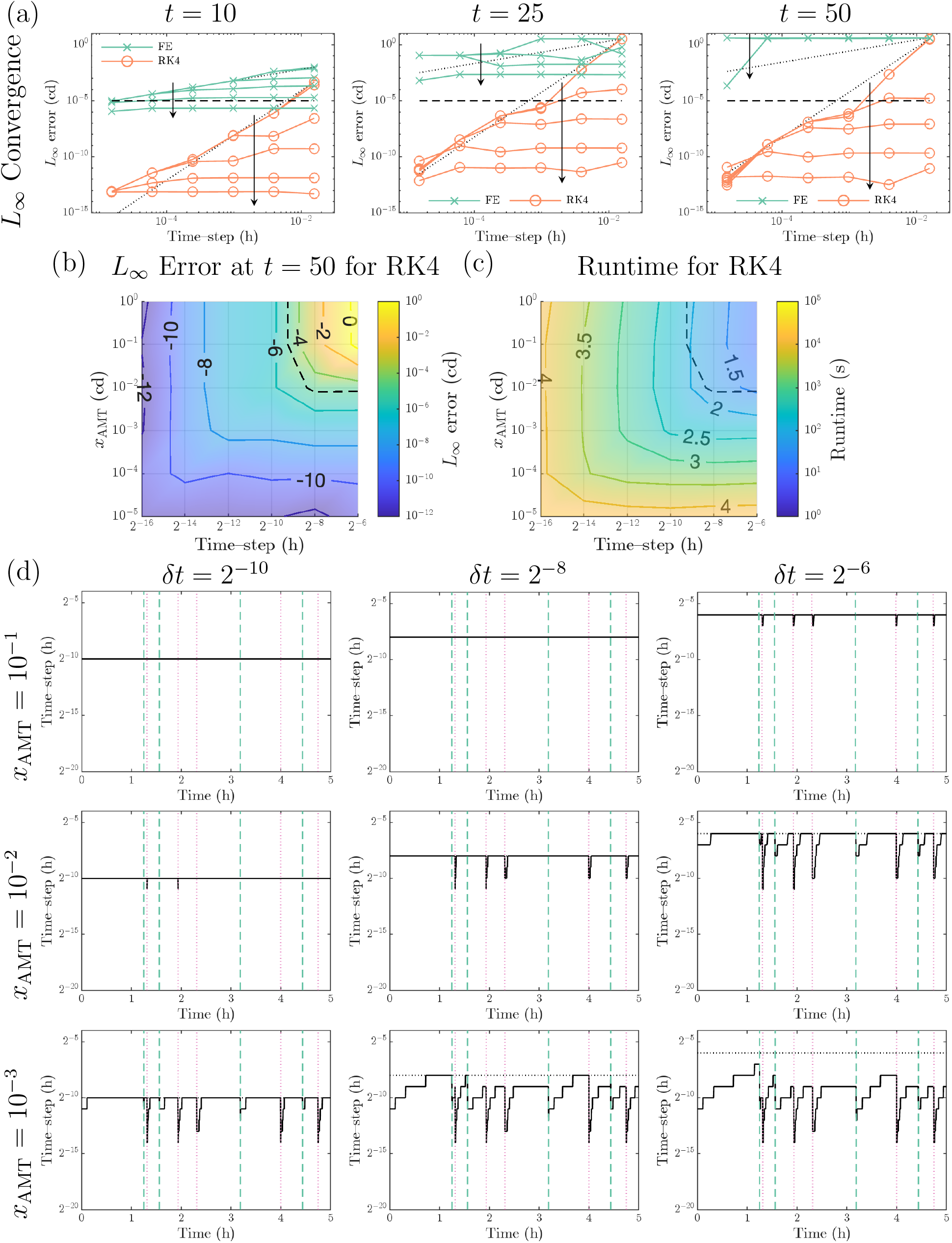
Adaptivity in simulations at dynamic equilibrium. Investigating how adaptivity inuences the convergence results from Figure 4. (a) how the L _∞_ error converges as *δt* is reduced for decreasing *x*_AMT_ at *t* = 10,25 and 50 for both FE and RK4 numerical methods. Arrow in direction of decreasing *x*_AMT_ from 1CD to 10^−5^CD. Dotted black lines show the theoretical order of convergence for each numerical method. (b) L_∞_ error for RK4 simulations as *δt* and *x*_AMT_ are varied. (c) Runtime for RK4 simulations as *δt* and *x*_AMT_ are varied. In (b) and contour labels show the exponent, *x*, of the *L*_∞_ error or runtime, 10^*x*^ The dashed contour in (b) and (c) is the target *L*_∞_ error of *L*_∞_ = 10^−5^. Similar plots for simulations using the FE numerical method show a constant *L*_∞_ error (except for the highest level of adaptivity) and are given in Supplementary Figure S3. (d) time–steps taken in simulations for various *δt* and *x*_AMT_. The time–step used by the adaptivity algorithm, 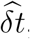, is given by the solid black line and the default time–step, *δt*, is given by the black dotted line (usually covered by the solid line). Red dotted lines are cell division events and green dot dashed lines show cell removal events (these are the same in all simulations).

Illustrations of time–steps taken are presented in Figure 5 (d). We see that the time–steps are reduced when events (births or deaths shown in red and green) happen. We also see that there is a balance to be made between global time–step, *δt* and adaptivity threshold, *x*_AMT_.

By looking at the contour *L*_∞_ = 10^−5^ (dashed black line in Figures 5 (b) and (c)) we can see that using RK4 we can get a the same *L*_∞_ error for a reduced runtime by using larger *δt* and smaller *x*_AMT_. To make this clearer we plot a magnified version of the runtime surface in Figure 6 (a). The ability to have an adaptive simulation with lower runtime and same *L*_∞_ error as a non adaptive simulation is because the *L*_∞_ = 10^−5^ contour (dashed line) in Figure 6 (a) intersects the *T*_R_ = 10^2.25^ seconds contour close to where there is no adaptivity and intersects the *T*_R_ = 10^1.5^ seconds contour where there is a larger time–step with adaptivity, these points are illustrated by the red and blue circles respectively in Figure 6 (a).

**Figure 6.**
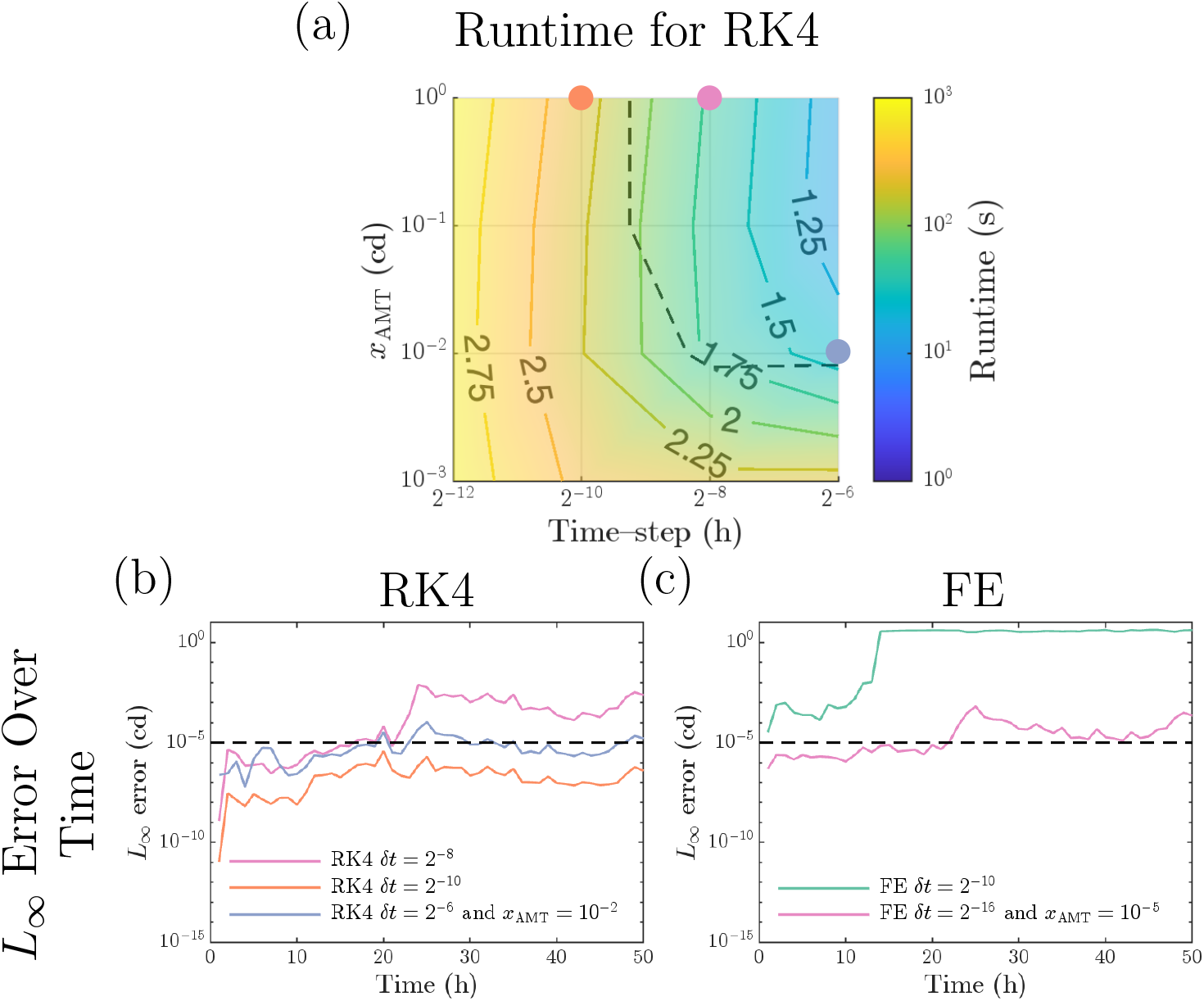
Adaptivity vs refinement. (a) Magnified version of simulation runtime from Figure 1 (b) highlighting region near target *L*_∞_ error contour (*L*_∞_ = 10). (b) *L*_∞_ error over time for numerical methods near target *L* error contour. Line colour matches the circled points in (a). The run time for these simulations is as follows: RK4 with *δt* = 2^−10^219 seconds; RK4 with *δt* = 2^−8^, 52 seconds; and RK4 with *δt* = 2^−6^ and *x*_AMT_ = 10^−2^, 24 seconds. (c) *L* error over time for simulations using Forward Euler with a similar run time to the simulations using RK4 in (b) *δt* = 2^−10^, 60 seconds (green) and one with comparable *L*_∞_ error FE with *δt* = 2 and *x*_AMT_ = 10^−5^, 6106 seconds (pink). Line colour matches the circled points in Supplementary Figure S3. The dashed contour in (a) and the dashed line in (b) and (c) denotes the target *L*_∞_ error (*L*_∞_ = 10^−5^).

In Figures 6 (b) and (c) we present the *L*_∞_ error over time for the numerical parameters above, and some similar choices. In (b) we show the *L*_∞_ error over time for RK4 simulations with: *δt* = 2^−8^; *δt* = 2^−10^; and *δt* = 2^−6^, *x*_AMT_ = 10^−2^. In (c) we show the *L*_∞_ error over time for a FE simulation of a similar runtime (*δt* = 2^−10^ green line) and a FE simulation with similar *L*_∞_ error to the optimal adaptive RK4 simulation (*δt* = 2^−16^ and *x*_AMT_ = 10^−5^, pink line). *L*_∞_ error and runtimes for these examples are given in Table 4.

**Table 4:**
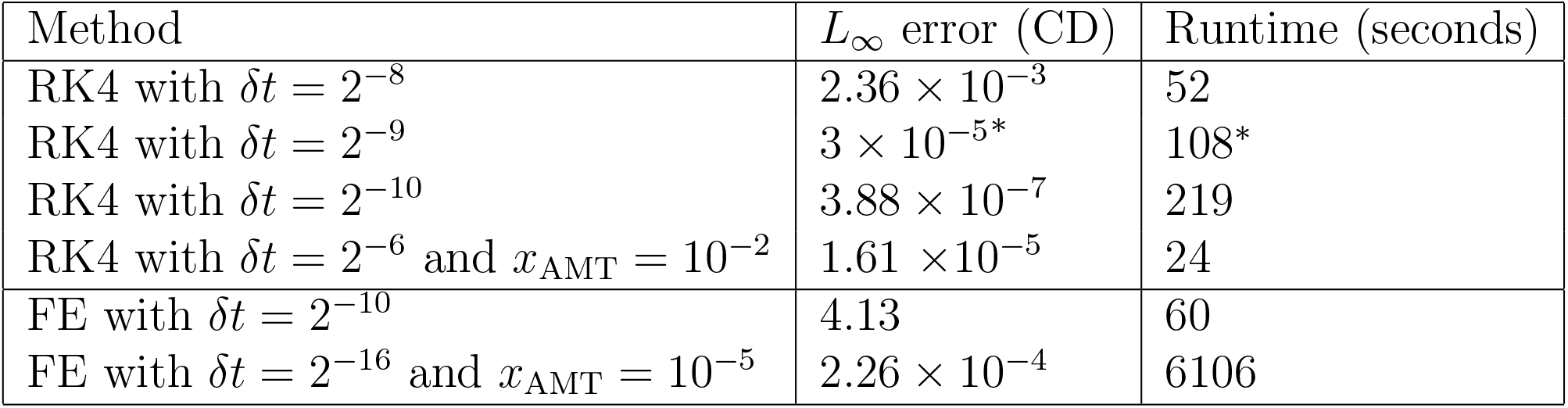
Error vs runtime with adaptivity. *L*_∞_ error and runtime for a selection of simulations from Figure 6. ^*^Denotes values calculated from interpolation from Figures 5 (b) and (c).

| Method | $L_\infty$ error (CD) | Runtime (seconds) |
| --- | --- | --- |
| RK4 with $\delta t = 2^{-8}$ | $2.36 \times 10^{-3}$ | 52 |
| RK4 with $\delta t = 2^{-9}$ | $3 \times 10^{-5*}$ | 108* |
| RK4 with $\delta t = 2^{-10}$ | $3.88 \times 10^{-7}$ | 219 |
| RK4 with $\delta t = 2^{-6}$ and $x_{\text{AMT}} = 10^{-2}$ | $1.61 \times 10^{-5}$ | 24 |
| FE with $\delta t = 2^{-10}$ | 4.13 | 60 |
| FE with $\delta t = 2^{-16}$ and $x_{\text{AMT}} = 10^{-5}$ | $2.26 \times 10^{-4}$ | 6106 |

To get any convergence at all with the FE method we require *δt* = 2^−16^ and *x*_AMT_ = 10^−5^ which takes 6106 seconds to run. We can achieve an equivalent *L*_∞_ error by running a RK4 simulation with *δt* = 2^−9^ which takes 108 seconds, a speed up of by a factor of 60. Moreover, an *L*_∞_ error of *L*_∞_ ≈ 10^−5^ can be achieved using RK4 either without adaptivity using *δt* = 2^−9^ (taking 108 seconds) or using *δt* = 2^−6^ and adaptivity with *x*_AMT_ = 10^−2^ (taking 24 seconds). A speed up by a factor of 4.

## 4 Conclusions

In this paper we have shown that by careful implementation of higher order numerical methods and by restricting events (births, deaths and “re–meshing”) to occur on common time steps, theoretical convergence of one step numerical methods, within multi–cellular simulations, can be demonstrated. Specifically the Runge–Kutta 4 method can be shown to have 4th order convergence. While implicit methods, such as Backward Euler, can be used their reliance on non–linear solvers causes any increase in stability to come at a large computational cost limiting their effectiveness.

In a growing system the Runge–Kutta 4 method can achieve the same accuracy over ten times faster than the FE method by being able to use a larger time–step, *δt*. For a system in dynamic equilibrium adaptivity can allow the Forward Euler method to maintain convergence. However, the Runge–Kutta 4 method (without adaptivity, with an appropriately chosen *δt*) can achieve the same accuracy 60 times faster. Adaptivity and a larger time–step can further reduce runtime of multicellular simulations using the Runge–Kutta 4 method by a factor of 4 leading to an over 200 fold reduction overall. While this work has demonstrated a significant speed up in multi–cellular simulations there are multiple avenues in which we could extend the work.

The current adaptive scheme only allows time–steps in multiples of 1*/*2, this assumption could be relaxed to have a more dynamic time–step calculation. Additionally, here we have focused on using a uniform time–step for all of the tissue. In the 3D Organ exemplar (much like for many multicellular simulations in the literature) we have a region of proliferative cells and region of differentiated cells. Where cells are proliferating we would need a small time–step, however, in the region where there is no (or even reduced) proliferation we could use a larger time–step. We could also only apply adaptivity (or different movement thresholds) in certain spatial regions to further reduce runtime without compromising on accuracy or stability. The selected regions would identify a set of cell connections (and therefore ODEs) which would require refined time–steps. Solutions from regions would be combined at common time–step as used in the purely temporal adaptive algorithm presented here.

The techniques presented here could also be extended to work with more detailed multi– cellular models, for example the rod based modelling framework [5] or immersed boundary method [42]. Here there would be issues with including virtual moves into higher order numerical methods and with coupling numerical methods for simulating cell movement with those for fluid flow.

In this paper we have demonstrated that the “default” numerical method of Forward Euler, while simple to implement, is not the best choice for multi–cellular models. While using a Runge–Kutta 4 scheme will take approximately 4 times longer (assuming that force evaluation is the dominant component of the simulation) for the same time–step, the improved accuracy will enable a much larger time–step to be used leading to a greater than tenfold reduction in overall runtime. Moreover, by using a simple adaptivity algorithm a larger global time–step can be used, with time–step refinement where required, the same accuracy can be achieved while reducing runtime, for our example simulation by a factor of 4.

The adaptive numerical framework presented here will enable existing simulations to be undertaken faster reducing the computational burden on researchers. In addition it will allow larger more detailed tissue simulations to be developed and simulated.

## Data availability

All code to reproduce this study is freely available, under an open source licence, from https://github.com/jmosborne/MultiCellularNumericalMethods.

## Acknowledgements

JMO would like to thank Dr Kathryn Tunyasuvunakool for her role in the initiation of this project as she laid the foundations for this work as part of her PhD project. JMO would also like to thank Kathryn’s co-supervisors Professor David Gavaghan, Professor Hillel Kugler, Dr Sara-Jane Dunn, and Professor Ruth Baker for their support of this project. JMO’s research is supported by the ARC (DP230100380, FT230100352).

## Supplementary Material

## Appendix A. Supplementary Figures

**Supplementary Figure 1:**
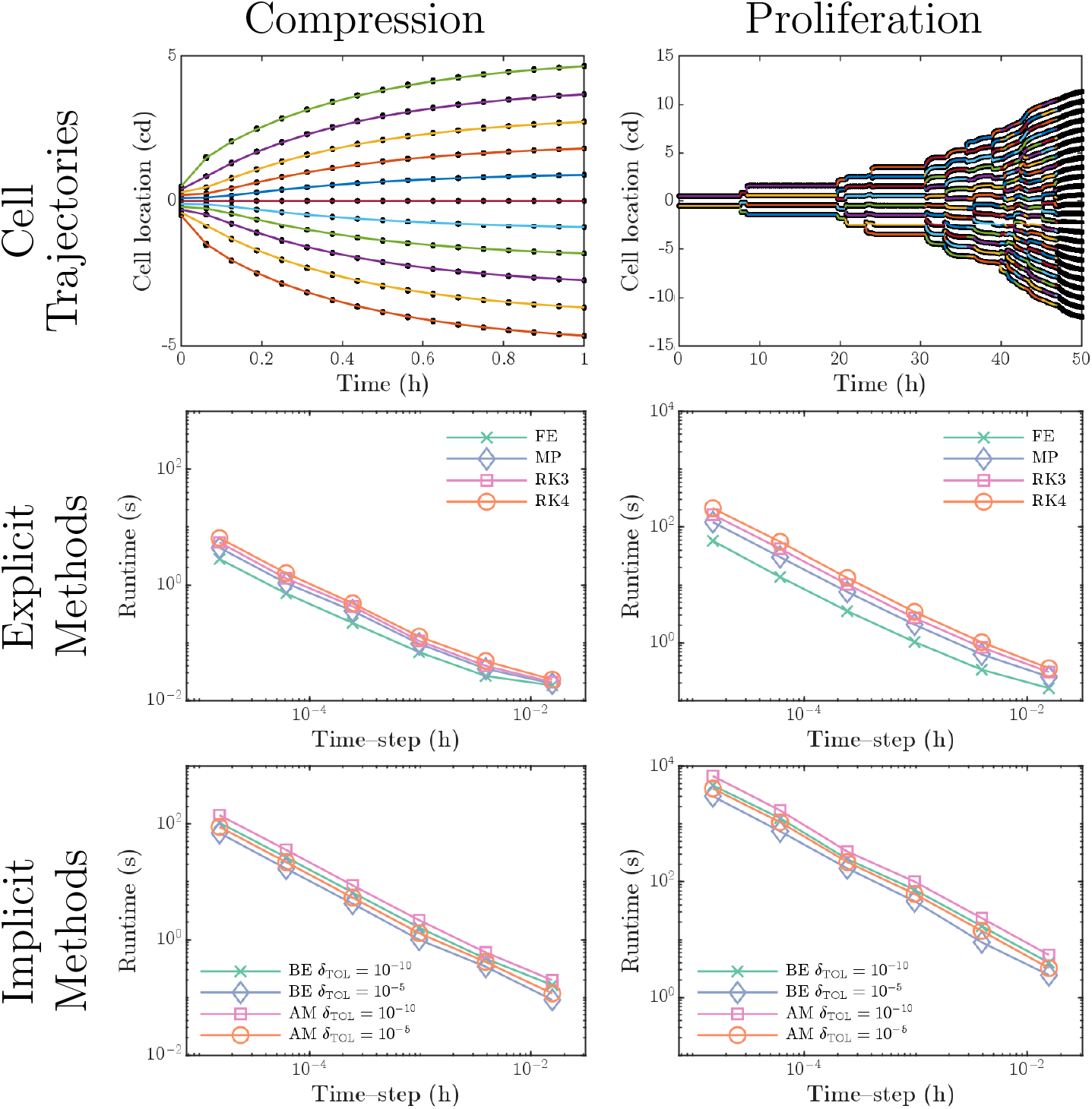
Runtime for one dimensional simulations. Runtimes for Supplementary Figure S1: **Cell trajectories and runtime for one dimensional simulations**. Top left, position of cells over time for 1D compression example. Coloured lines represent the exact solution and the numerical solutions at 2^−4^ hour intervals are given by by the black dots. Top right, position of cells over time, coloured lines represent the exact solution and the numerical solutions at 1 hour intervals are given by the black dots. Middle and bottom, simulation runtime for simulations from Figure 2

**Supplementary Figure 2:**
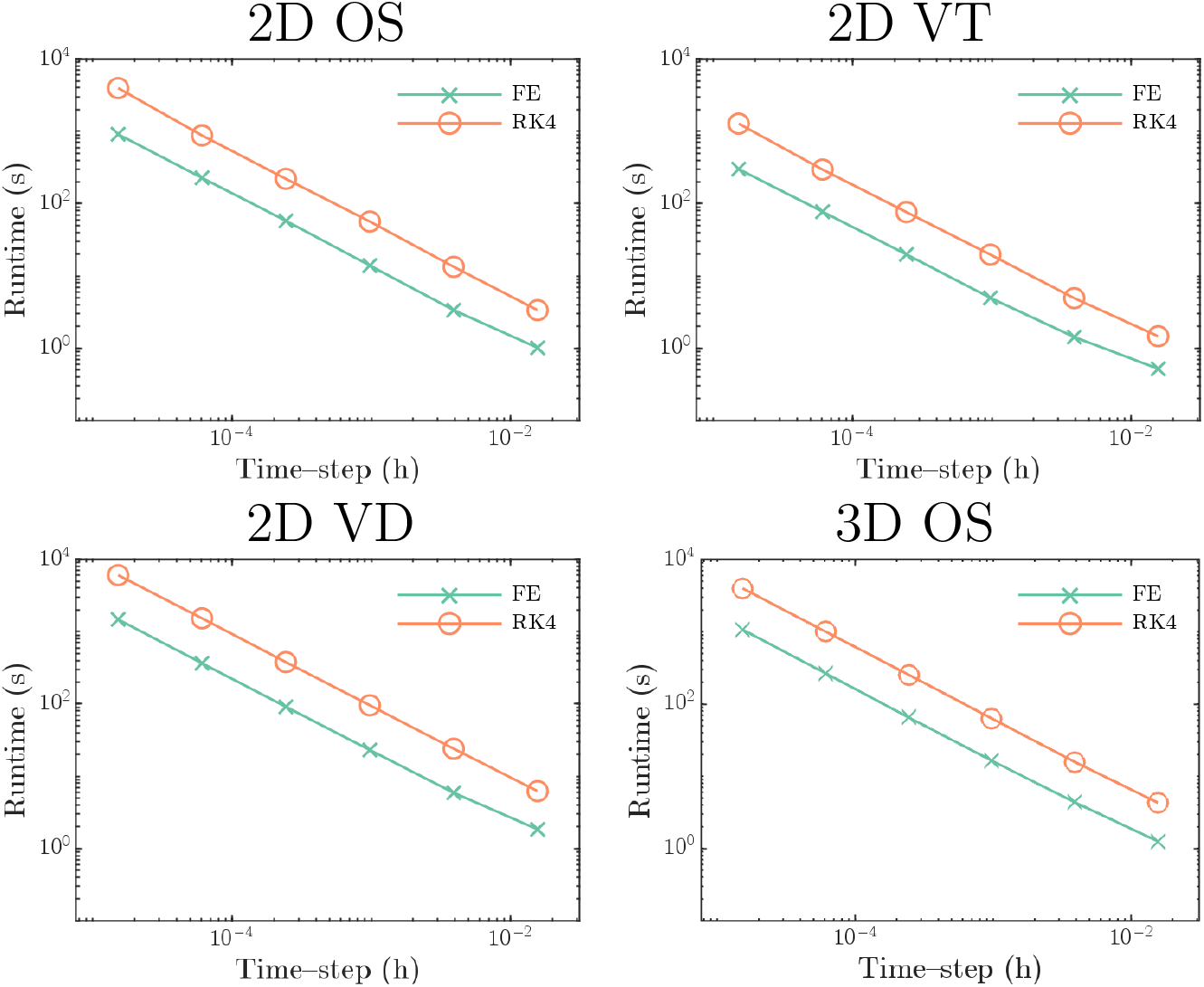
Runtime for two and three dimensional simulations. Runtimes as *δt* is varied for simulations using FE and RK4 methods from Figure 3.

**Supplementary Figure S3:**
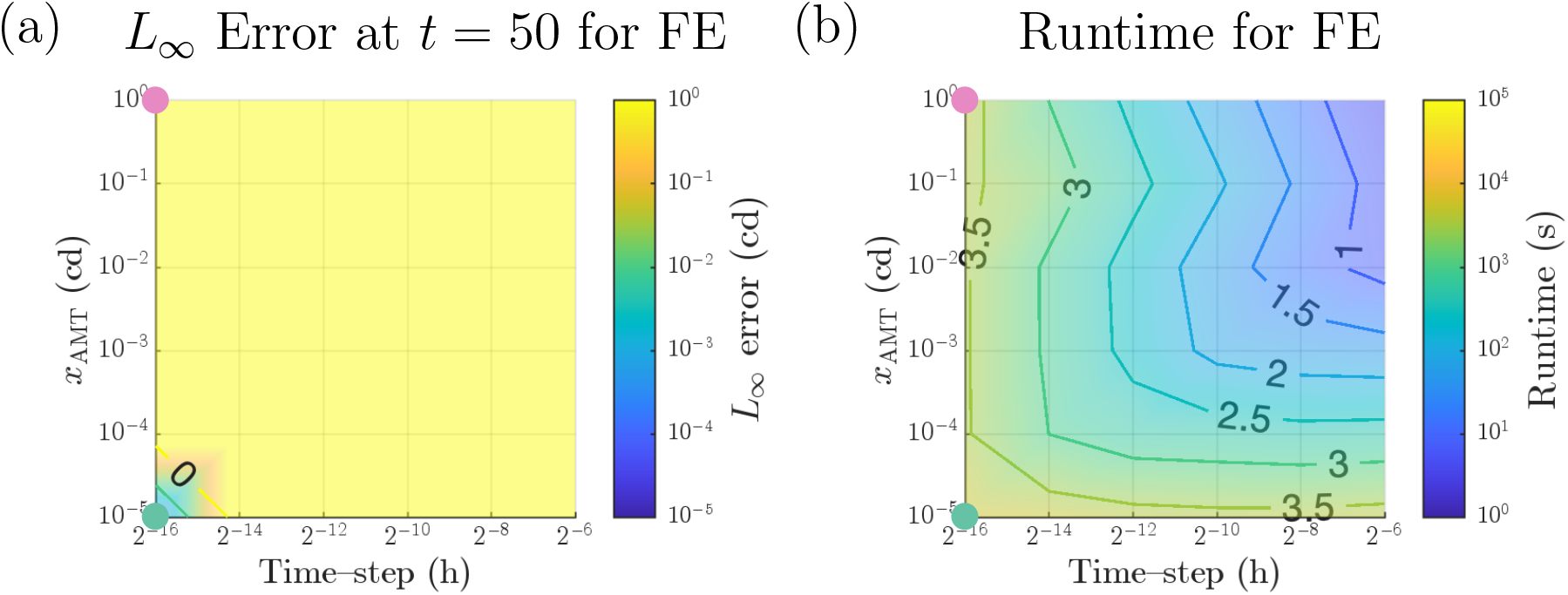
Adaptivity using FE in simulation of dynamic equilibrium. (a) *L* _*∞*_ error as *δt* and *x*_AMT_ are varied. (b) *L* error as *δt* and *x*_*AMT*_ are varied. varied. Contour labels show the exponent, *x*, of the *L*_∞_ error or runtime, 10^*x*^. Green and pink circles represent the numerical parameters of the simulations in Figure 6 (c).

## Appendix B. Analytic solution of 1D Compression and Proliferation exemplars

For a 1D chain of cells using a linear spring model it is possible to find an exact solution for the motion of the cells. In this section we detail this solution and show how to use it to get exact solutions for comparison for the 1D Compression and 1D Proliferation exemplars. For the 1D Compression exemplar the equations of motion, for cell centres *x*_0_, …, *x*_*n*_ are

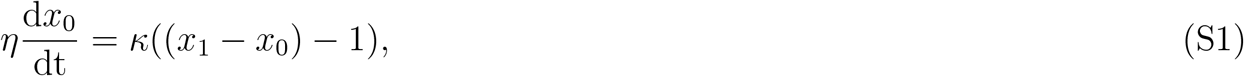

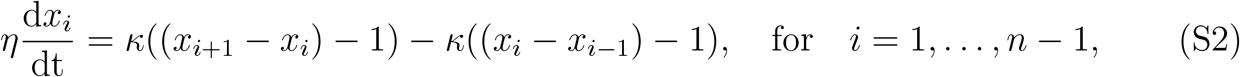

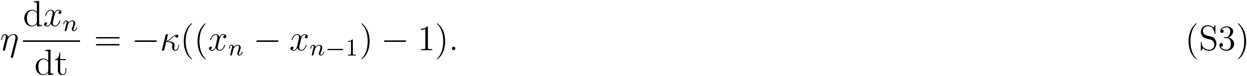

Where *η* is the drag coefficient for the cell centres and *κ* is the spring coefficient representing the mechanical interaction between cell centres. We can rearrange these equations as

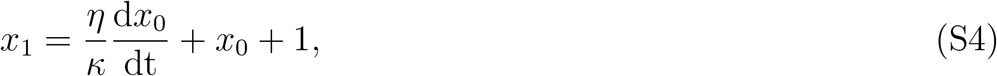

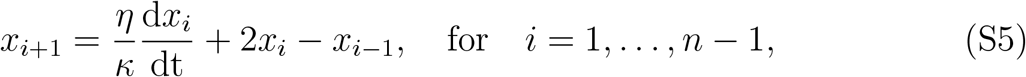

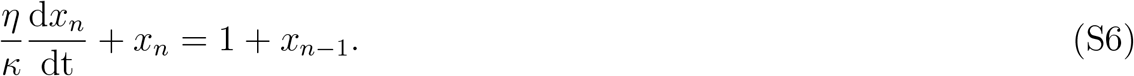

Equations (S4) and (S5) allow the calculation of *x*_1_, …, *x*_*n*_ given some *x*_0_ and then Equation (S6) gives a further condition on the solutions. Therefore we look for a solution of the form

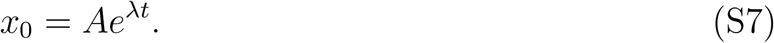

Substituting Equation (S7) into Equation (S4) and then Equation (S5) allows us to calculate expressions for *x*_1_, …, *x*_*n*_

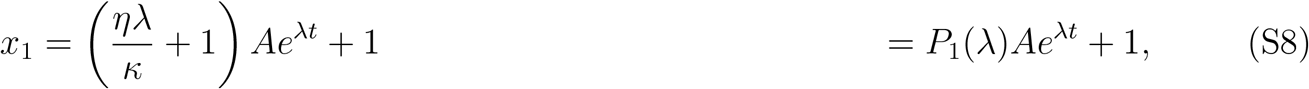

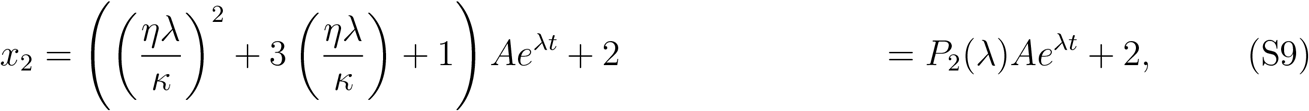

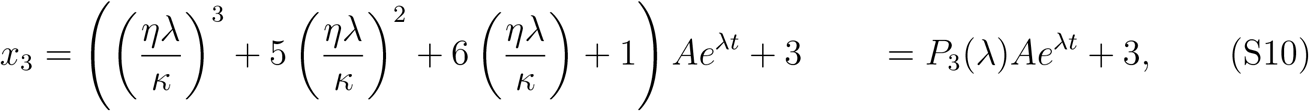

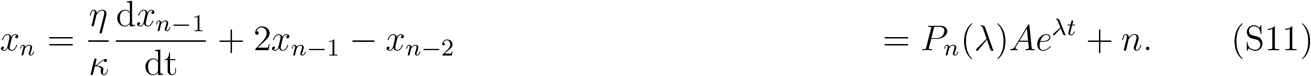

Where *P*_*i*_(*λ*) is an *i*th degree polynomial, defined and calculated as above. We then substitute Equation (S11) into Equation (S6) to give

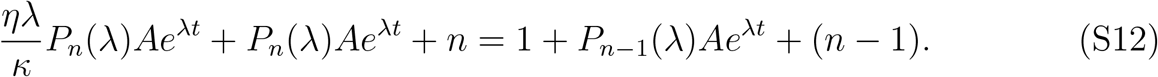

Which can be rearranged to

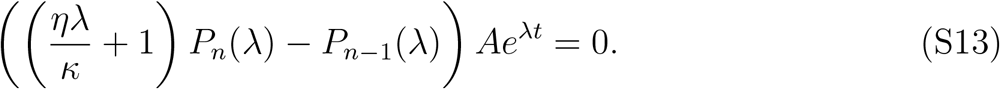

So to avoid having a trivial solution, *x*_*i*_ = *i* (i.e., no displacement from equilibrium), we need

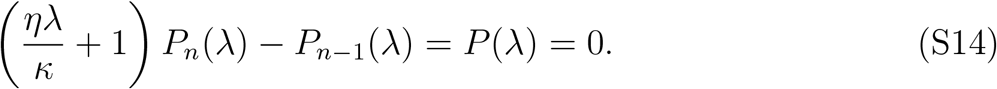

Where *P* (*λ*) is the *n* + 1st degree characteristic polynomial of the system where the roots *λ*_*i*_ give non trivial solutions. Therefore the solution, *x*_0_, is of the form

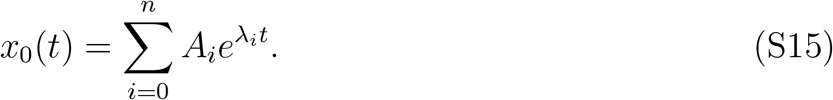

Where *λ*_*i*_ are the roots of Equation (S14) and the coefficients *A*_*i*_ are chosen to satisfy the initial conditions 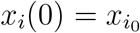. We can combine Equation (S15) with Equations (S8)–(S11), to derive the solution for *x*_*i*_ for *i* = 1, …, *n*,

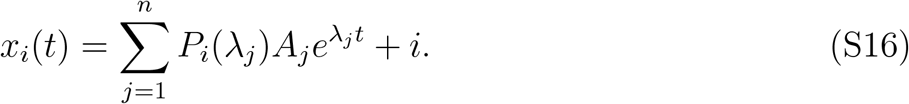

Using this we can set up the following linear system

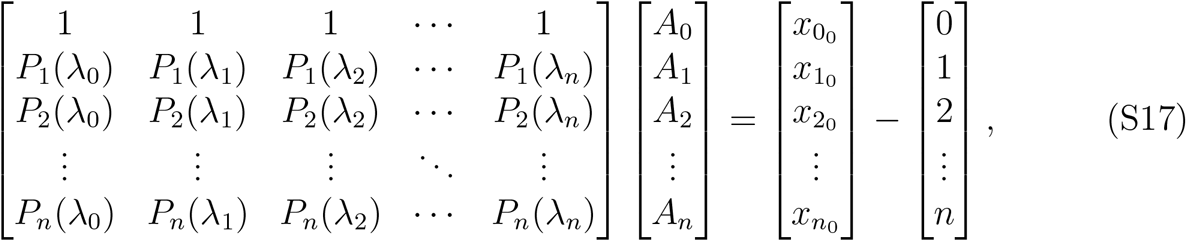

which can be solved for the coefficients *A*_*i*_. This gives us the analytic solutions to the 1D Compression problem for any initial placement of cells. An example of this analytic solutions is shown in Supplementary Figure S1 top left.

The analytic solution of the 1D Proliferation example can also be calculated by specifying the division locations and using the solution method above in between division events. An example of this analytic solution with divisions is shown in Supplementary Figure S1 top right.

## Footnotes

1 It is not possible to calculate the exact solution in these cases as the models are intractable to analytic methods in more than one dimension.

